# Generating high-quality libraries for DIA-MS with empirically-corrected peptide predictions

**DOI:** 10.1101/682245

**Authors:** Brian C. Searle, Kristian E. Swearingen, Christopher A. Barnes, Tobias Schmidt, Siegfried Gessulat, Bernhard Kuster, Mathias Wilhelm

**Affiliations:** Institute for Systems Biology, Seattle, WA, USA; Proteome Software, Inc. Portland, OR, USA; Novo Nordisk Research Center Seattle, Inc. Seattle, WA, USA; Chair of Proteomics and Bioanalytics, Technical University of Munich, Freising, Germany; SAP SE, Potsdam, Germany; Bavarian Center for Biomolecular Mass Spectrometry, Freising, Germany

## Abstract

Data-independent acquisition approaches typically rely on sample-specific spectrum libraries requiring offline fractionation and tens to hundreds of injections. We demonstrate a new library generation workflow that leverages fragmentation and retention time prediction to build libraries containing every peptide in a proteome, and then refines those libraries with empirical data. Our method specifically enables rapid library generation for non-model organisms, which we demonstrate using the malaria parasite *Plasmodium falciparum*, and non-canonical databases, which we show by detecting missense variants in HeLa.

## INTRODUCTION

Data-independent acquisition (DIA) mass spectrometry (MS) is a powerful label-free technique for deep, proteome-wide profiling.^1,2^ With DIA, mass spectrometers are tuned to systematically acquire tandem mass spectra at regular retention time and m/z intervals, making the method free of intensity-triggering biases introduced by data-dependent acquisition (DDA). To accomplish this, precursor isolation windows are widened such that multiple peptides are usually co-fragmented in the same MS/MS scan. DIA methods generally identify peptides with library search engines^3–5^ using sample-specific spectrum libraries^6^ from DDA experiments. In peptide-centric searching,^7^ library entries are scored according to retention time, such that the best-scoring time point for each peptide is reported. Only peptides present in the libraries can be detected, and the peptide detection reports must be corrected to limit the number of potential false discoveries.^8^ Most importantly, these libraries are built at the expense of time, sample, and considerable effort with offline fractionation, especially considering that they are typically not reusable across laboratories or instrument platforms.^9^

When sample-specific spectrum library generation is either impossible or impractical, as is frequently the case with non-model organisms, sequence variants, splice isoforms, or scarce sample quantities, software tools such as Pecan^10^ and DIA-Umpire^11^ can detect peptides from DIA experiments without a spectrum library by directly searching every peptide in FASTA databases. Gas-phase fractionation^12^ (GPF) improves detection rates with these tools^10^ by injecting the same sample multiple times with tiled precursor isolation windows, allowing each injection to have narrower windows (and thus fewer co-fragmented peptides) with the same instrument duty cycle. While this method is often prohibitively expensive because it requires enough instrument time and protein content for multiple injections for each sample, the use of multiple GPF injections can be applied just to pooled samples to generate DIA-only chromatogram libraries that make it easier to detect peptides in single-injection DIA experiments.^13^ However, even when using GPF these tools still generally detect fewer peptides than library search engines, which can leverage previously acquired instrument-specific fragmentation and measured retention times. Recently it has become possible to accurately predict spectra from peptide sequences,^14,15^ but direct searching of single-injection DIA data has remained problematic, in part due to the false discovery rate (FDR) correction required when considering all possible tryptic peptides in a FASTA database. Proteins show 3-4 orders of magnitude difference in intensity between the best- and worst-responding tryptic peptides,^16^ and only considering the best-responding peptides in libraries can improve detection rates by lessening the required FDR correction.

## RESULTS

### Generating empirically-corrected libraries from peptide predictions

Here we report on a DIA-only workflow that produces higher-quality libraries than those generated by DDA while simultaneously supplanting the need for any offline fractionation. Our workflow (Fig. 1a, Online Methods) uses a recently developed deep neural network, Prosit,^14^ to predict a library of fragmentation patterns and retention times for every peptide in FASTA databases. Fragmentation prediction in Prosit adjusts based on normalized collision energy (NCE), and we tune the NCE parameter for each peptide charge state to account for DIA-specific fragmentation. Building on the chromatogram library method,^13^ we collect 6 GPF-DIA runs of a sample pool using 90-minute gradients (12 total hours of MS acquisition). We then search these GPF-DIA runs against the predicted library using EncyclopeDIA to detect peptides in the pool. Assuming that virtually every consistently quantifiable peptide in an experiment is detectable from the pool using GPF-DIA, we filter the predicted library to remove peptides and charge states that cannot be found in the pool. We also replace the retention times and fragmentation patterns in the predicted library with sample- and instrument-specific values found in the 6 runs. The resulting empirically-corrected library typically contains only 10-15% of the entries in the predicted library, requiring less stringent FDR correction when later searching single-injection DIA runs.

**Figure 1:**
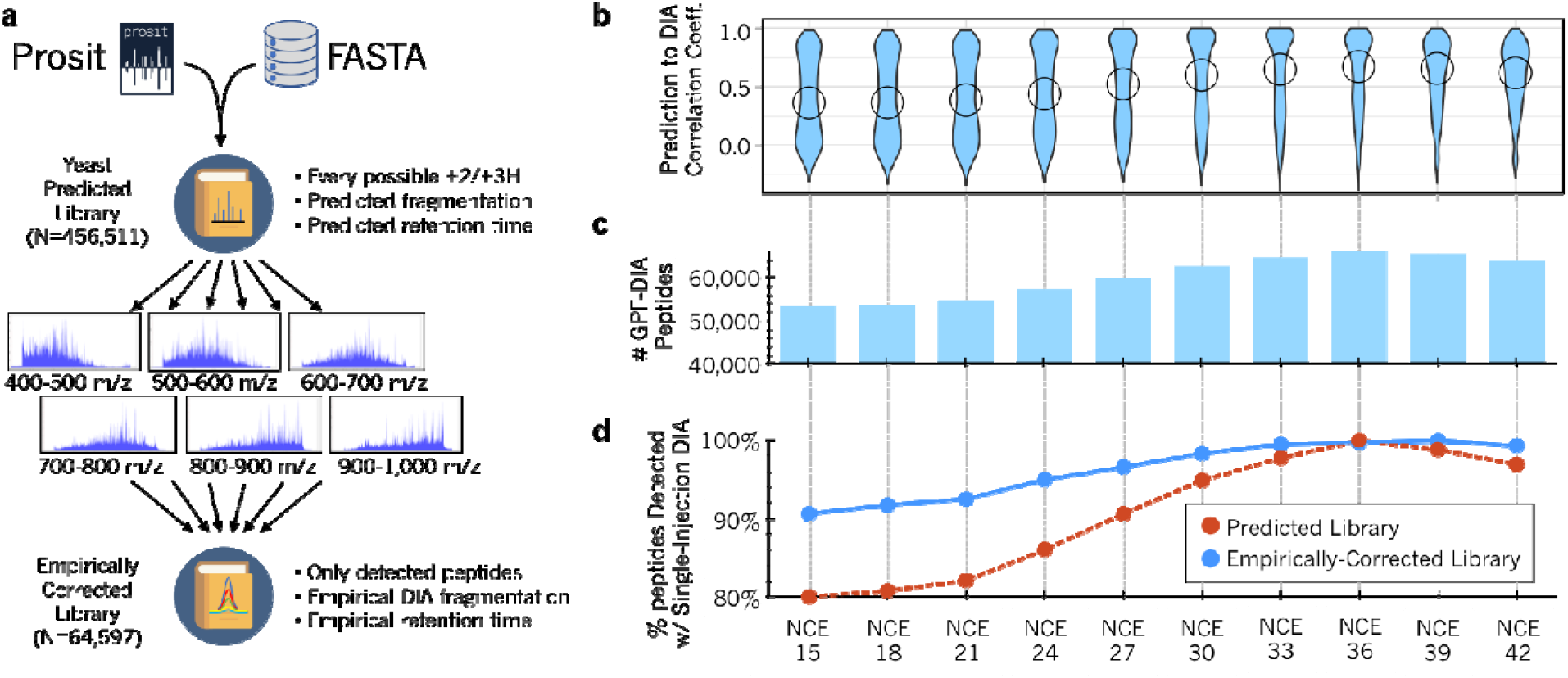
Evaluating empirically-corrected libraries made with peptide predictions. (**a**) Workflow for generating empirically-corrected libraries from Prosit predictions of peptides in FASTA databases. GPF-DIA runs of a sample pool are searched with a spectrum library of fragmentation and retention time predictions. Peptides detected in these runs are compiled with fragmentation patterns and retention times extracted from the GPF-DIA data into an empirically-corrected library. (**b**) Violin plots showing spectral correlation between predicted library for yeast peptides and single-injection DIA at various NCE settings (circles indicate medians). (**c**) The total numbers of empirically-corrected library entries detected from GPF-DIA at various NCE settings. (**d**) The fraction of peptides detected in single-injection DIA relative to the optimal NCE for empirically-corrected libraries (blue line) are more consistent than predicted libraries (red dashed line) across a wide range of NCE settings.

### Validating the empirically-corrected library methodology

Although searching single-injection DIA runs directly with predicted libraries is highly dependent on prediction accuracy, we found that our workflow produced high-quality libraries even if the predicted libraries were not precisely tuned for the instrument, suggesting broad cross-platform applicability. We demonstrated this by modulating the NCE setting in Prosit (Fig. 1b) and found that with a wide range of NCE settings, searching GPF-DIA spectra of yeast acquired on a Fusion Lumos MS produced between 33% and 60% larger empirically-corrected libraries than a sample-specific 10-fraction DDA library containing 40,025 unique peptides (Fig. 1c). Additionally, searches were less sensitive to prediction accuracy after empirical correction (Fig. 1d), in part due to consistently improved fragmentation patterns (Supp. Fig. 1).

While offline fractionation requires additional liquid chromatography (LC) using orthogonal separation modes to online LC-MS, GPF occurs completely within the mass spectrometer, making it both more reproducible and easier to perform. Also unlike offline fractionation, GPF also maintains the same LC matrix complexity as each experimental sample, ensuring more precise retention time consistency. This process produced libraries with better retention-time accuracy (80% of peptides within 35 seconds) than both the predicted (80% within 5.4 minutes) and DDA libraries (80% within 4.6 minutes), even when the DDA libraries were acquired on the same instrument (Supp. Fig. 2). Coupled with smaller library size and improved fragmentation patterns, these 3 factors had roughly equal and orthogonal improvements over directly searching single-injection DIA with predicted libraries (Supp. Fig. 3). Overall, searching single-injection DIA runs with empirically corrected libraries detected 66% more yeast peptides than searching the 10-fraction DDA library, and 37% more peptides than a 39-injection DDA library in reanalyzed HeLa datasets^13^ from a Q-Exactive HF (QE-HF) MS (Fig. 2a).

**Figure 2:**
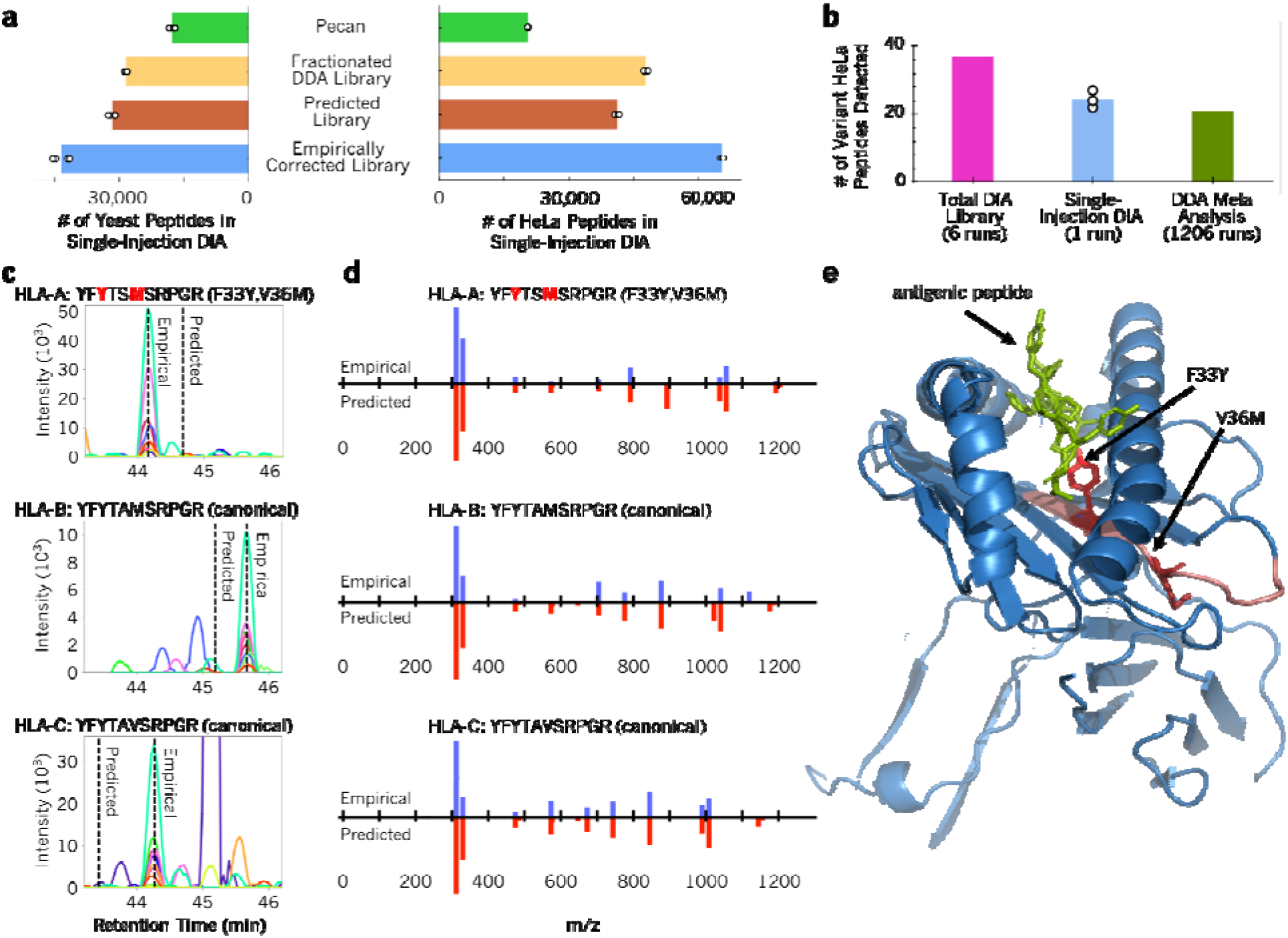
Empirically-corrected libraries improve peptide and missense variant detection rates. (**a**) The average number of yeast (N=4) and HeLa (N=3) peptides in single-injection DIA detected using library-free searching with Pecan and searching either a fractionated DDA spectrum library, a predicted spectrum library, or an empirically-corrected library. (**b**) The number of HeLa missense variants detected in the total DIA library, single-injection DIA (triplicate experiments), or a meta-analysis of 40 published DDA experiments,^22^ each filtered to a 1% peptide/protein FDR. (**c**) Retention time predictions differ somewhat from empirical data from GPF-DIA for homologous peptides: YFYTSMSRPGR from HLA-A (F33Y,V36M), YFYTAMSRPGR from HLA-B, and YFYTAVSRPGR from HLA-C. (**d**) Relative +1H y-type and b-type fragmentation patterns above 200 m/z for the same peptides are shown as butterfly plots with empirical intensities (blue) above predicted intensities (red). (**e**) All 3 peptides are in the HLA peptide binding/presentation region, as indicated by YFYTSMSRPGR (red) in HLA-A (blue) relative to an antigenic peptide (green).

### Finding missense sequence variants with non-canonical libraries

Most publicly available spectrum libraries^17–19^ are built by searching millions of spectra against a canonical genome. Our approach facilitates the analysis of sequence variants and splice junctions by simplifying exome-specific library construction, and we demonstrate this by analyzing HeLa-specific missense variants. RNA-seq expression across different HeLa strains^20^ suggests that 12.4k genes containing 127 missense variants determined by the COSMIC^21^ cell line project are typically expressed (>=10 counts, >=0.1 RPKM). Recently a custom database including COSMIC variants was used to reanalyze 40 public HeLa datasets containing 1,206 total DDA runs,^22^ globally detecting 21 missense variants. In contrast, from 6 GPF-DIA runs we built an empirically-corrected HeLa-specific library with 7,484 protein groups including 37 missense variants (Supp. Data 1), approximately 30% of the expected expressed variants reported by COSMIC. With the library we detected an average of 24 missense variants from single-injection DIA runs (Fig. 2b). While in general the additionally detected variant-containing peptides will not greatly affect the overall number of quantitative measurements, quantifying key peptides in specific genes, such as the peptide antigen binding region of HLA^23^ (Fig. 2c-e), can have a profound effect on biological interpretation. While improved fragmentation and retention time accuracy in empirically-corrected libraries can buffer search engines from over-reporting missense variants, care must be taken when attempting to distinguish between homologous sequences with DIA. Heterozygous peptides with similar retention times, such as those associated with G623S from the kinase EEF2K (Supp. Fig 4a), can share fragment ions (Supp. Fig 4b) and must be localized as if they contained post-translational modifications^24,25^ should they fall in the same precursor isolation window.

### Detecting and quantifying peptides from non-model organisms

We further applied our workflow to analyze cultures of *Plasmodium falciparum*, the parasite responsible for 50% of all malaria cases. We performed DDA and DIA on late-stage (stage IV/V) gametocytes, the form of the parasite that is transmitted from an infected human to the mosquito vector that spreads the disease. Using a QE-HF MS, we produced a 3,415-protein empirically corrected library, increasing the previously known proteome^26^ size by 58% (Fig. 3a) while only missing 2.5% of proteins detected with other methods. Using this library we measured 2,740 proteins in single-injection DIA experiments of *P. falciparum* peptides. Because *Plasmodium* is an obligate parasite, even samples produced by *in vitro* culture are frequently contaminated with the host proteome. To study the robustness of our method we diluted *P. falciparum* peptides into peptides derived from uninfected red blood cells, and found we could detect >80% of the parasite peptides in up to 1:99 dilution (Fig. 3b, Supp. Fig. 5a). We found that with our method, DIA maintained higher quantification precision and depth than comparable DDA experiments at each dilution step (Fig. 3c, Supp. Fig. 5b), which were analyzed with MaxQuant^27^ with “match between runs” enabled. Leveraging the highly accurate retention times in our empirically-corrected library, we found that we could in part recover contaminated *P. falciparum* samples (Supp. Fig. 6), even when MaxQuant/DDA reported the majority of peptides as missing values.

**Figure 3:**
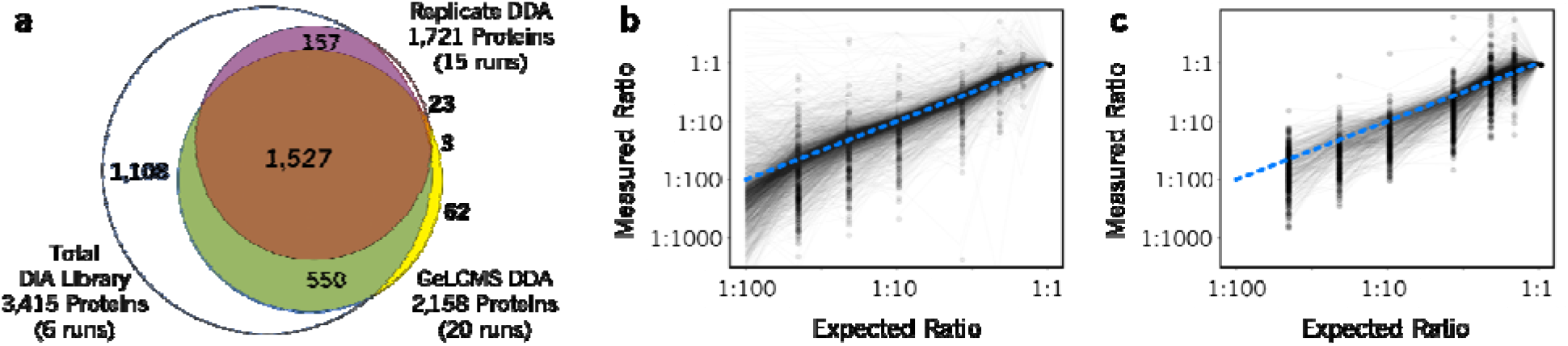
Rapid library generation and quantitation of *Plasmodium falciparum* proteins. **(a)** A proportionally-sized Venn diagram showing the overlap between proteins detected in the empirically-corrected library (blue), the full single-injection dilution curve with DDA (red), and the Stage IV/V mixed male/female sexual *P. falciparum* proteome published by Lasonder *et al.*^*26*^ from 20x GeLCMS-fractionated DDA runs (yellow). Quantitative ratios of (**b**) 2,740 DIA- and (**c**) 1,491 DDA-detected malaria parasite *P. falciparum* proteins (gray lines) after dilution with uninfected RBC lysate relative to the undiluted sample (N=1). Dots indicate the lowest dilution level of measurement for each protein, and the blue dashed line indicates the ideal measurement. Empirically-corrected libraries enabled DIA quantification of 2,152 *P. falciparum* proteins in the last dilution sample (1:99), even when no proteins could be detected or quantified using DDA.

We analyzed 2,444 *P. falciparum* protein groups with at least 2 peptides (Supp. Data 2) that were detected in every DIA run of 3 culture replicates with 3 technical replicates each (9 runs total). After cross-referencing against the PlasmoDB^28^ compendium of 20 different studies at various asexual, sexual, and mosquito stages, we observed 20 proteins previously undetected by mass spectometry in any life stage of *P. falciparum*. We found an additional 396 proteins that had never been observed in late-stage gametocytes, including 55 previously detected only in immature (stage I/II) gametocytes. Experiments such as these have the potential to redefine the protein expression signature for each stage of the *Plasmodium* life cycle, information that may be vital to identifying targets to disrupt parasite maturation. For example, among those proteins previously undetected in gametocytes is the calcium-dependent protein kinase CDPK2. *P. falciparum* parasites lacking CDPK2 develop normally through asexual stages, but male gametocytes are incapable of undergoing exflagellation to become gametes, thereby preventing transmission to the mosquito vector.^29^ Our work validates that CDPK2 is indeed present at measurable levels in gametocytes, paving the way to monitor dynamic expression of this kinase over the course of parasite maturation.

## DISCUSSION

In conclusion, empirical correction of predicted spectrum libraries enables rapid library generation for non-canonical proteomes or non-model organisms without offline fractionation. To encourage the reuse of this method, we have released a growing repository of pre-generated predicted libraries compatible with EncyclopeDIA, Skyline, and Scaffold DIA, which are available for download at ProteomicsDB (https://www.proteomicsdb.org/prosit/libraries). We also developed a graphical user interface in EncyclopeDIA (Supplementary Note) to facilitate making empirically-corrected libraries for new proteomes from any FASTA database.

## Data availability statement

Prosit (https://www.proteomicsdb.org/prosit) and EncyclopeDIA 1.0 (https://bitbucket.org/searleb/encyclopedia) are both available under the Apache 2 open-source license. The raw data from the yeast and *P. falciparum* studies are available at MassIVE (MSV000084000). The raw data from the HeLa reanalysis are available as originally published at MassIVE (MSV000082805).

## Acknowledgements

We thank S. Kappe for insightful discussions, N. Carmago for providing the gametocyte samples, L. Pino for providing yeast lysates, B. Kim for technical assistance.We also thank M. MacCoss, K. Grove, and M. Guldbrandt for providing instrument time. B.C.S. is supported by the Translational Research Fellows Program (TRFP) from the Institute for Systems Biology. K.E.S. is supported by K25 AI119229. This work was in part funded by the German Federal Ministry of Education and Research (BMBF, grant no. 031L0008A and no. 031L0168) and the EU Horizon 2020 grant EPIC-XS (grant no. 823839).

## Author contributions

B.C.S. conceived the study. B.C.S., K.E.S., and C.A.B. performed the experiments. B.C.S., T.S., and S.G. developed the software. B.C.S., B.K., and M.W. supervised the work. All authors wrote and approved the manuscript.

## METHODS

### *Saccharomyces cerevisiae* culture and sample preparation

As described in Searle *et al.*,^13^ *S. cerevisiae* strain BY4741 (Dharmacon) was cultured at 30°C in YEPD and harvested at mid-log phase. Cells were pelleted and lysed in a buffer of 8 M urea, 50 mM Tris (pH 8), 75 mM NaCl, 1 mM EDTA (pH 8) followed by 7 cycles of 4 minutes bead beating with glass beads. After a 1 minute rest on ice, lysate was collected by piercing the tube and centrifuging for one minute at 3,000 × g and 4°C into an empty eppendorf. After further centrifugation at 21,000 × g and 4°C for 15 minutes, the protein content of the supernatant was removed and estimated using BCA. Proteins were then reduced with 5 mM dithiothreitol for 30 minutes at 55°C, alkylated with 10 mM iodoacetamide in the dark for 30 minutes at room temperature, and diluted to 1.8 M urea, before digestion with sequencing grade trypsin (Pierce) at a 1:50 enzyme to substrate ratio for 16 hours at 37°C. 5N HCl was added to approximately pH 2 to quench the digestion, and the resulting peptides were desalted with 30 mg MCX cartridges (Waters). Peptides were dried with vacuum centrifugation and brought to 1 μg / 3 μl in 0.1% formic acid (buffer A) prior to mass spectrometry acquisition.

### *Plasmodium falciparum* culture and red blood cell sample preparation

Three flasks of stage IV/V *P. falciparum* NF54 gametocytes were prepared. Asexual cultures were synchronized with sorbitol and set up at 5% hematocrit and 1% parasitemia. Gametocytogenesis was induced by withholding fresh blood and allowing parasitemia to increase. N-acetyl glucosamine was added to media for 4 days beginning 7 days after set-up in order to remove asexual parasites. Gametocyte-infected erythrocytes (giRBC) were enriched from uninfected erythrocytes (uiRBC) by magnetic-activated cell sorting at stage III. Stage IV/V gametocytes were collected on day 15 post-set-up. Additional uiRBC were also prepared by washing multiple times in RPMI and stored at 50% hematocrit. giRBC and uiRBC cells were lysed in a buffer of 10% sodium dodecyl sulfate (SDS), 100 mM ammonium bicarbonate (ABC), cOmplete EDTA-free Protease Inhibitor Cocktail (Sigma), and Halt Phosphatase Inhibitor Cocktail (Thermo Scientific). Proteins were then reduced with 20-40 mM tris(2-carboxyethyl)phosphine (TCEP) for 10 minutes at 95°C and alkylated with 40-80 mM iodoacetamide in the dark for 20 minutes at room temperature. After centrifugation at 16,000 × g to pellet insoluble material, proteins were purified with methanol:chloroform extraction^30^ and dried and resuspended in 8M urea buffer before the content was estimated using BCA. After dilution to 1.8 M urea, proteins were digested with sequencing-grade trypsin (Promega) at a 1:40 enzyme-to-substrate ratio for 15 hours at 37°C. The resulting peptides were desalted with Sep-Pak cartridges (Waters), dried with vacuum centrifugation, and brought to 1 μg / 3 μl in 0.1% formic acid (buffer A) prior to mass spectrometry acquisition. In addition, several digested peptide mixtures were made by diluting peptides from one flask of giRBC cells with peptides from uiRBC cells at ratios of 1:0, 2:1, 7:8, 4:15, 1:9, 2:41, 2:91, and 1:99 giRBC:uiRBC.

### Liquid chromatography mass spectrometry (*S. cerevisiae*)

Tryptic *S. cerevisiae* peptides were separated with a Thermo Easy nLC 1200 on self-packed 30 cm columns packed with 1.8 μm ReproSil-Pur C18 silica beads (Dr. Maisch) inside of a 75 μm inner diameter fused silica capillary (#PF360 Self-Pack PicoFrit, New Objective). The 30 cm column was coiled inside of a Sonation PRSO-V1 column oven set to 35°C prior to ionization into the MS. The HPLC was performed using 200 nl/min flow with solvent A as 0.1% formic acid in water and solvent B as 0.1% formic acid in 80% acetonitrile. For each injection, 3 μl (approximately 1 μg) was loaded and eluted with a linear gradient from 7% to 38% buffer B over 90 minutes. Following the linear separation, the system was ramped up to 75% buffer B over 5 minutes and finally set to 100% buffer B for 15 minutes, which was followed by re-equilibration to 2% buffer B prior to the subsequent injection. Data were acquired using data-independent acquisition (DIA).

The Thermo Fusion Lumos was set to acquire 6 GPF-DIA acquisitions of a biological sample pool using 120,000 precursor resolution and 30,000 fragment resolution. The automatic gain control (AGC) target was set to 4e5, the maximum ion inject time (IIT) was set to 60 ms, the normalized collision energy (NCE) was set to 33, and +2H was assumed as the default charge state. The GPF-DIA acquisitions used 4 m/z precursor isolation windows in a staggered window pattern with optimized window placements (i.e., 398.4 to 502.5 m/z, 498.5 to 602.5 m/z, 598.5 to 702.6 m/z, 698.6 to 802.6 m/z, 798.6 to 902.7 m/z, and 898.7 to 1002.7 m/z). Individual samples for proteome profiling acquisitions used single-injection DIA acquisitions (120,000 precursor resolution, 15,000 fragment resolution, AGC target of 4e5, max IIT of 20 ms) using 8 m/z precursor isolation windows in a staggered window pattern with optimized window placements from 396.4 to 1004.7 m/z.

For generation of an *S. cerevisiae* spectral library, 80 μg of the same tryptic digests described above were separated into 10 total fractions using the Pierce high-pH reversed-phase peptide fractionation Kit (Thermo, #84868). Briefly, peptides were loaded onto hydrophobic resin spin column and eluted using the following 8 acetonitrile steps: 5%, 7.5%, 10%, 12.5%, 15%, 17.5%, 20.0%, and 50%, keeping both the wash and flow-through. The resulting peptide fractions were injected into the same Thermo Fusion Lumos using the same chromatography setup and column described above, but configured for data-dependent acquisition (DDA). After adjusting each fraction to an estimated 0.5 to 1.0 μg on column, the fractions were measured in a top-20 configuration with 30 second dynamic exclusion. Precursor spectra were collected from 300-1650 m/z at 120,000 resolution (AGC target of 4e5, max IIT of 50 ms). MS/MS were collected on +2H to +5H precursors achieving a minimum AGC of 2e3. MS/MS scans were collected at 30,000 resolution (AGC target of 1e5, max IIT of 50 ms) with an isolation width of 1.4 m/z with a NCE of 33.

### Liquid chromatography mass spectrometry (*P. falciparum*)

Tryptic *P. falciparum* and RBC peptides were separated with a Thermo Easy nLC 1000 and emitted into a Thermo Q-Exactive HF. In-house laser-pulled tip columns were created from 75 μm inner diameter fused silica capillary and packed with 3 μm ReproSil-Pur C18 beads (Dr. Maisch) to 30 cm. Trap columns were created from Kasil fritted 150 μm inner diameter fused silica capillary and packed with the same C18 beads to 2 cm. The HPLC was performed using 250 nl/min flow with solvent A as 0.1% formic acid in water and solvent B as 0.1% formic acid in 80% acetonitrile. For each injection, 3 μl (approximately 1 μg) was loaded and eluted using a 84-minute gradient from 6 to 40% buffer B, followed by steep 5-minute gradient from 40 to 75% buffer B and finally set to 100% buffer B for 15 minutes, which was followed by re-equilibration to 0% buffer B prior to the subsequent injection. Data were acquired using either DDA or DIA.

The Thermo Q-Exactive HF was set to acquire DDA in a top-20 configuration with “auto” dynamic exclusion. Precursor spectra were collected from 400-1600 m/z at 60,000 resolution (AGC target of 3e6, max IIT of 50 ms). MS/MS were collected on +2H to +5H precursors achieving a minimum AGC of 1e4. MS/MS scans were collected at 15,000 resolution (AGC target of 1e5, max IIT of 25 ms) with an isolation width of 1.4 m/z with a NCE of 27. Additionally, 6 GPF-DIA acquisitions were acquired of a biological sample pool (60,000 precursor resolution, 30,000 fragment resolution, AGC target of 1e6, max IIT of 60 ms, NCE of 27, +3H assumed charge state) using 4 m/z precursor isolation windows in a staggered window pattern with optimized window placements (i.e., 398.4 to 502.5 m/z, 498.5 to 602.5 m/z, 598.5 to 702.6 m/z, 698.6 to 802.6 m/z, 798.6 to 902.7 m/z, and 898.7 to 1002.7 m/z). Individual samples used single-injection DIA acquisitions (60,000 precursor resolution, 30,000 fragment resolution, AGC target of 1e6, max IIT of 60 ms) using 16 m/z precursor isolation windows in a staggered window pattern with optimized window placements from 392.4 to 1008.7 m/z.

### FASTA databases and predicted spectrum libraries

Species-specific reviewed FASTA databases for *Homo sapiens* (April 25 2019, 20415 entries) and *Saccharomyces cerevisiae* (January 25 2019, 6729 entries) were downloaded from Uniprot. The *Plasmodium falciparum* FASTA database^31^ was downloaded from PlasmoDB^28^ version 43 (April 24 2019, 5548 entries). The Ensembl-based HeLa-specific FASTA database^22^ was downloaded from the ACS Publications website and modified to be compatible with EncyclopeDIA (47305 entries, including both canonical and variant protein sequences). Each database was digested *in silico* to create all possible +2H and +3H peptides with precursor m/z within 396.43 and 1002.70, assuming up to one missed cleavage. Peptides were further limited to be between 7 and 30 amino acids to match the restrictions of the Prosit tool.^14^ In general, NCE were assumed to be 33 (yeast was processed using NCE from 15 to 42 in 3 NCE increments) but modified to account for charge state. Since DIA assumes all peptides are of a fixed charge, we adjusted the NCE setting as if peptides were fragmented at the wrong charge state using the formula:

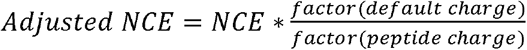

where the factors were 1.0 for +1H, 0.9 for +2H, 0.85 for +3H, 0.8 for +4H, and 0.75 for +5H and above.^32^ After submitting to Prosit, predicted MS/MS and retention times were converted to the EncyclopeDIA DLIB format for further processing. Scripts to produce Prosit input from FASTAs and build EncyclopeDIA-compatible spectrum libraries from Prosit output are available as functions in EncyclopeDIA 1.0.

### DDA data processing

All Thermo RAW files were converted to .mzML format using the ProteoWizard package^33^ (version 3.0.18299) using vendor peak picking. DDA data was searched with Comet^34^ (version 2017.01 rev. 1), allowing for fixed cysteine carbamidomethylation, variable peptide n-terminal pyro-glu, and variable protein n-terminal acetylation. Fully tryptic searches were performed with a 50 ppm precursor tolerance and a 0.02 Da fragment tolerance permitting up to 2 missed cleavages. High-pH reverse-phase fractions were combined and search results were filtered to a 1% peptide-level FDR using PeptideProphet^35^ from the Trans-Proteomic Pipeline^36^ (TPP version 5.1.0). A yeast-specific Bibliospec^37^ DDA spectrum library was created from Thermo Q-Exactive DDA data using Skyline^38^ (Daily version 19.0.9.149).

*P. falciparum* and RBC DDA data was additionally processed with MaxQuant^27^ (version 1.6.5.0) to perform label-free quantitation with precursor ion integration. MaxQuant was configured to use default parameters, briefly fixed cysteine carbamidomethylation, variable methionine oxidation, and variable protein n-terminal acetylation. Fully tryptic searches were performed with a 20 ppm fragment tolerance using both the human and *P. falciparum* FASTA databases, as well as common contaminants and filtered to a 1% peptide-level FDR. Quantification was performed using unique and razor peptides with the match between runs setting turned on.

### DIA data processing

DIA data was overlap demultiplexed^39^ with 10 ppm accuracy after peak picking in ProteoWizard (version 3.0.18299). Searches were performed using EncyclopeDIA (version 0.8.3), which was configured to use default settings: 10 ppm precursor, fragment, and library tolerances. EncyclopeDIA was allowed to consider both B and Y ions and trypsin digestion was assumed. EncyclopeDIA search results were quantified and filtered to a 1% peptide-level using Percolator^40^ (version 3.1) and 1% protein-level FDR assuming parsimony.

### Empirically-corrected library generation

Predicted libraries were corrected with EncyclopeDIA using the chromatogram library method described previously.^13^ Briefly, GPF-DIA samples for a given study were loaded into EncyclopeDIA using the above parameters, where the search library was set to the appropriate predicted spectrum library. Peptides detected by EncyclopeDIA were exported as a chromatogram library.

A peptide entry in a chromatogram library is similar to a peptide entry in a spectrum library in that it contains a precursor mass, retention time, and a fragmentation spectrum. In addition, a chromatogram library entry also contains the peptide peak shape, and a correlation score for every fragment ion indicating the agreement between the overall peptide peak shape and the fragment peak shape. This score provides an indication of the likelihood the fragment ion was interfered with in the GPF-DIA, with the expectation that it will also likely be interfered with in single-injection DIA as well. Missense variants in the HeLa empirically-corrected library were manually validated.

This process created a new empirically-corrected library containing only peptides found in the GPF-DIA samples, and also retained empirical fragment ion intensities and retention times observed from the DIA data. These libraries were made to be compatible with both EncyclopeDIA and Skyline and were used for downstream analysis of single-injection DIA.

