## Supplementary Information for "Generating high-quality libraries for DIA-MS with empirically-corrected peptide predictions"

**SUPPLEMENTARY NOTE**

**Using EncyclopeDIA and Prosit to Make Empirically-Corrected Peptide Libraries**

Data-independent acquisition (DIA) mass spectrometry is a powerful label-free proteomics technique. DIA methods typically rely on sample-specific spectrum libraries from deeply fractionated data-dependent acquisition (DDA) experiments. This tutorial (**based on EncyclopeDIA version 0.8.3**) will walk you through how to create equal- or higher-quality libraries using only DIA data.

**Prerequisites**

Prosit is a web application that is accessible from any operating system without installation. EncyclopeDIA is a cross-platform Java application that has been tested for Windows, Macintosh, and Linux. EncyclopeDIA requires 64-bit Java 1.8 or higher. If you don’t already have it, you can download “Windows x64” from:

http://www.oracle.com/technetwork/java/javase/downloads/jre8-downloads-2133155.html

Alternatively, you can also use the open-source OpenJDK if the Oracle’s Java license is too restrictive. After you have 64-bit Java 1.8, double click on the EncyclopeDIA .JAR file to launch the GUI interface. If you are using a Macintosh, you may need to right click on the EncyclopeDIA .JAR and select “Open” to execute it for the first time with the proper permissions.

**Collecting DIA Data**

We recommend consulting this general-purpose overview for collecting DIA data: <https://bitbucket.org/searleb/encyclopedia/downloads/dia%20methods%20setup.pdf>

We recommend these parameters for collecting single-injection and gas-phase fractionated DIA:

<https://docs.google.com/spreadsheets/d/1A8AQlmLroAkQcAcsiGTNvnGBE2lGpkMwhh0YLTBHXKA>

**Building Prosit CSV Input with EncyclopeDIA**

The “Convert/Create Prosit CSV from FASTA” menu option launches a dialog for building a Prosit CSV. Select the input FASTA, charge range, the number of maximum missed cleavages, and the m/z range of interest. In general, because FDR is calculated based on the total number of peptides searched, we recommend using small FASTA databases (fewer than 25,000 entries) and narrow charge ranges to shrink the necessary search space:

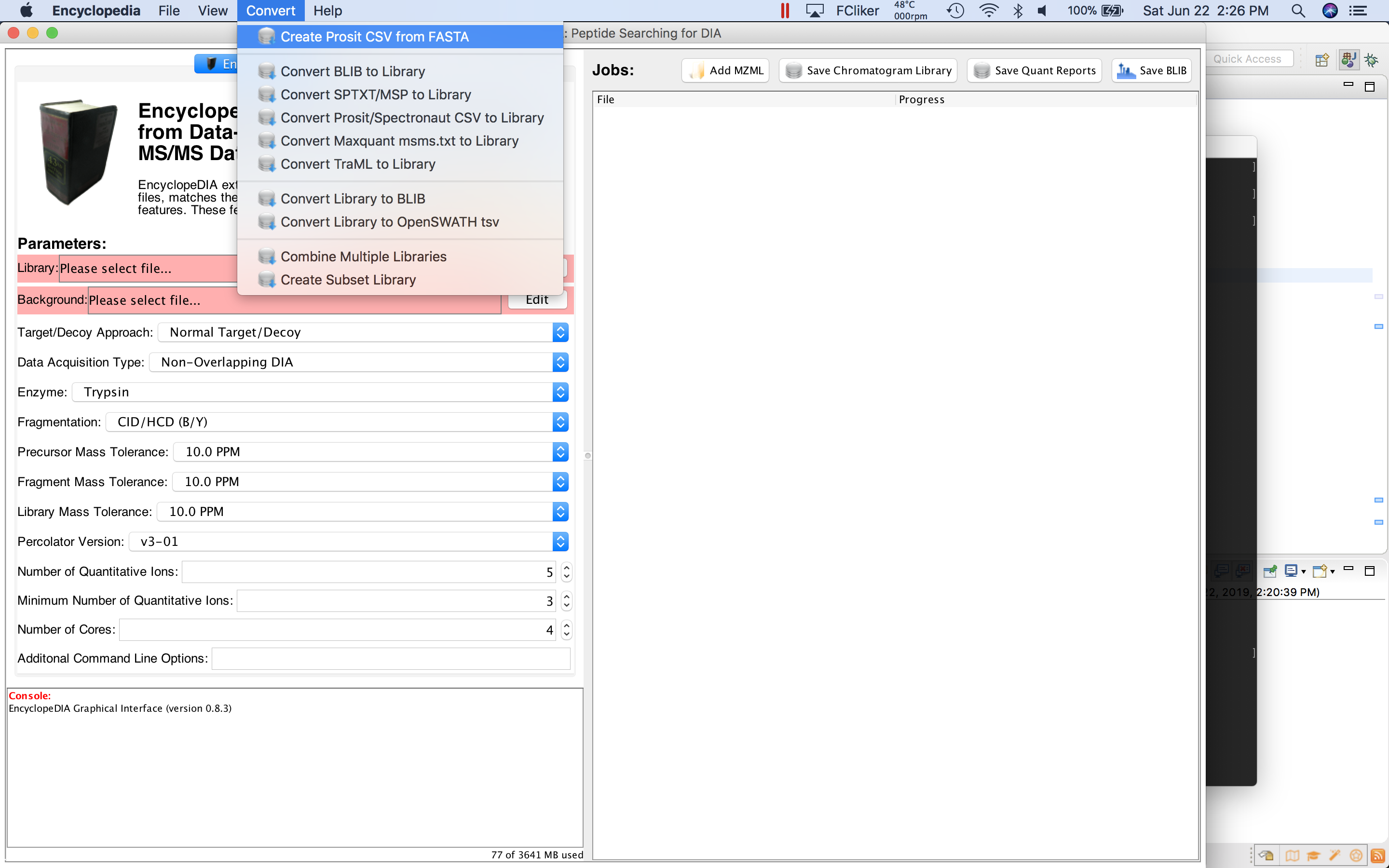

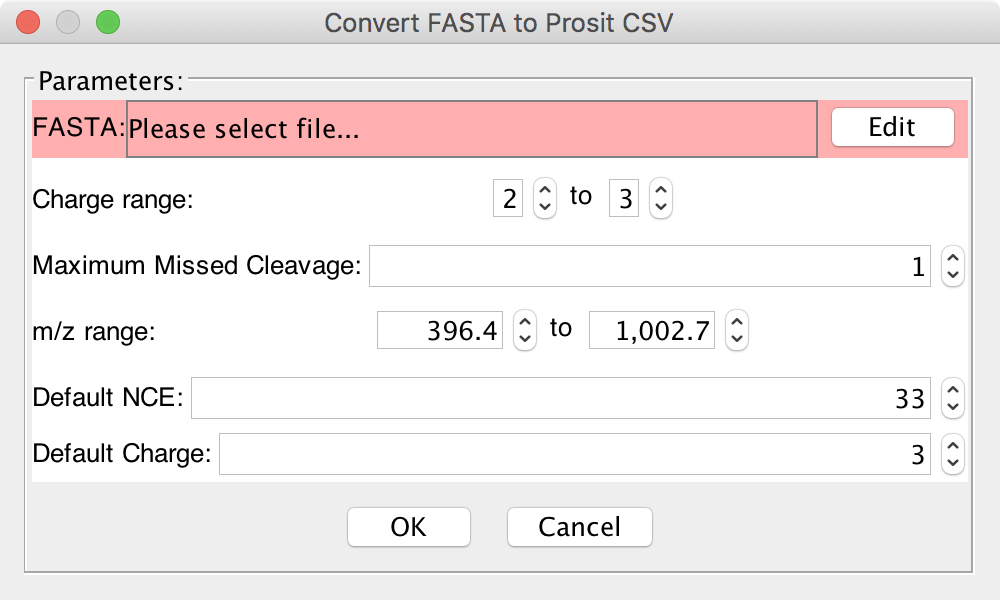

Based on how you acquired your DIA experiment, set the target NCE and default charge state for the prediction. With DIA all peptides are fragmented assuming they are the same charge. For peptides that are not the default charge state, EncyclopeDIA back calculates what the Prosit NCE should be. NCE settings are based on the Thermo Fusion Lumos mass spectrometer. The default (33 NCE) should work in most cases, but if you’d like to fine-tune this setting then the following conversions are a good starting point:

- Fusion-class orbitrap instruments: Actual NCE setting
- Q-Exactive-class orbitrap instruments: Actual NCE setting + 6
- ToF instruments: EV/2.5

This will create a CSV file in the same directory as your FASTA.

**Generating Prosit Predictions**

First, go to the Prosit website at <https://www.proteomicsdb.org/prosit/>. Using the “Spectral Library” tab and follow the three steps:

1. “Settings”: select “CSV” to provide the list of peptides, then hit “next”
2. “Upload Files”: click the cloud icon to upload the CSV from EncyclopeDIA, then hit “next”
3. “Task ID”: select “Generic text” for the return format and hit “submit”

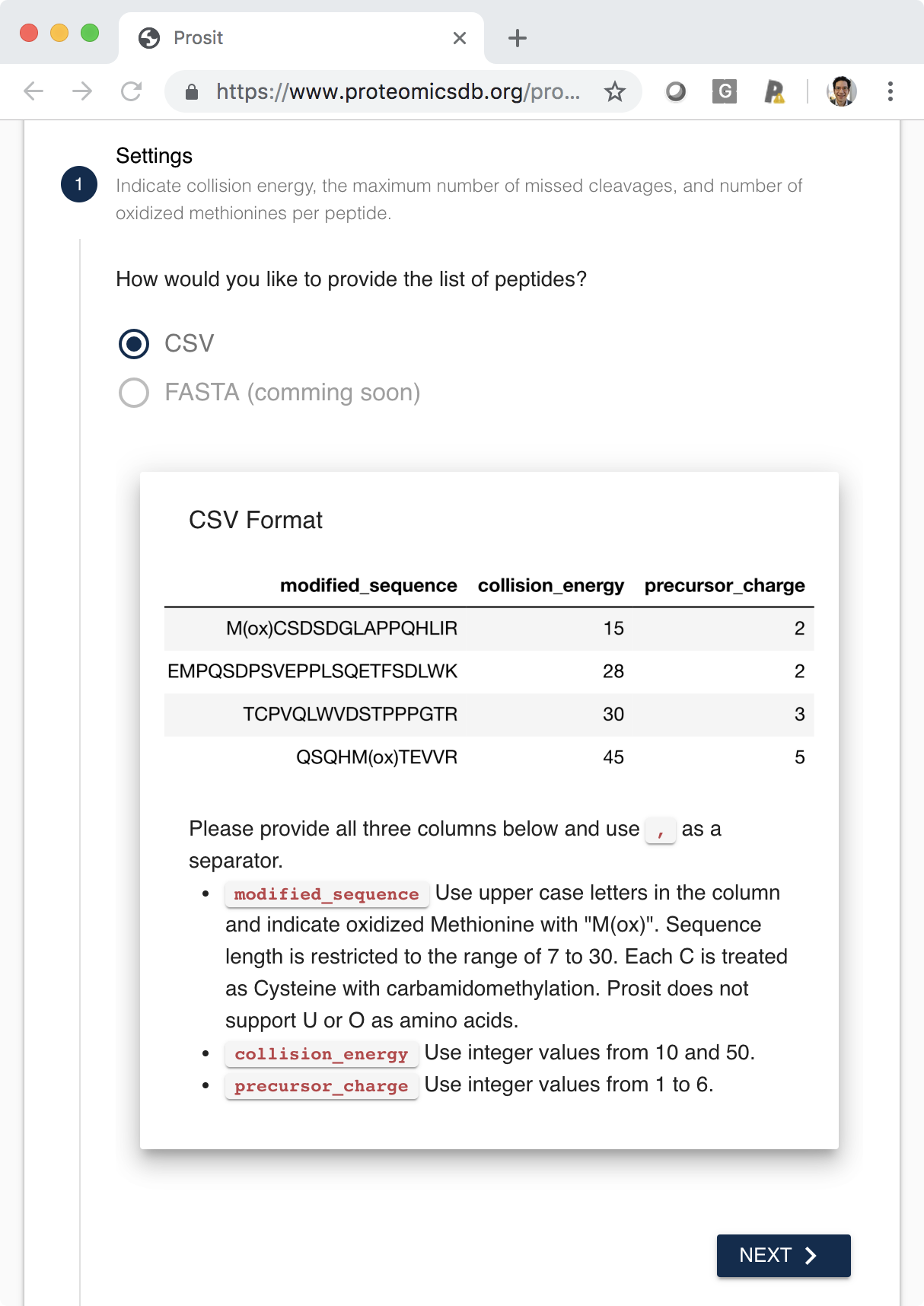

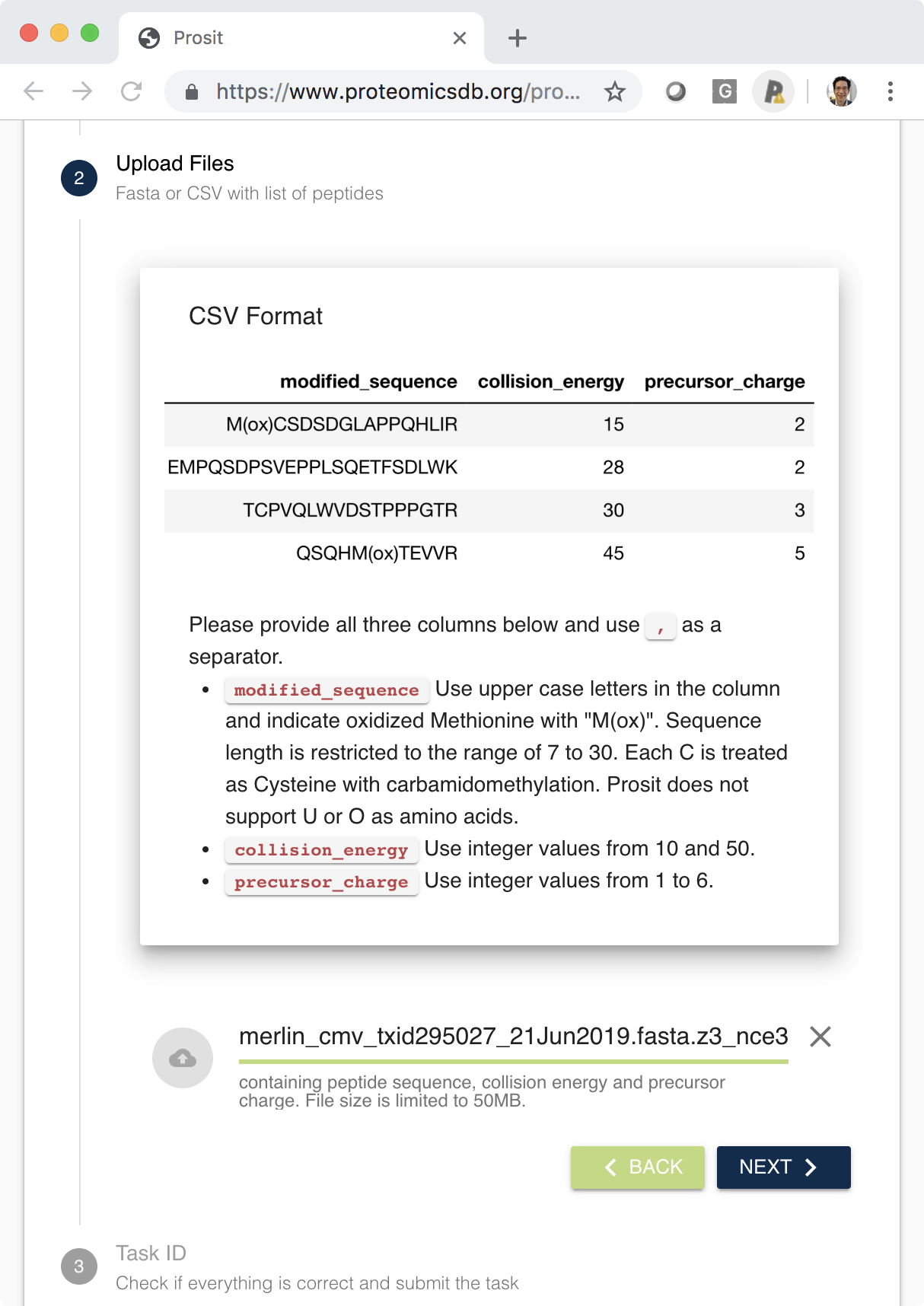

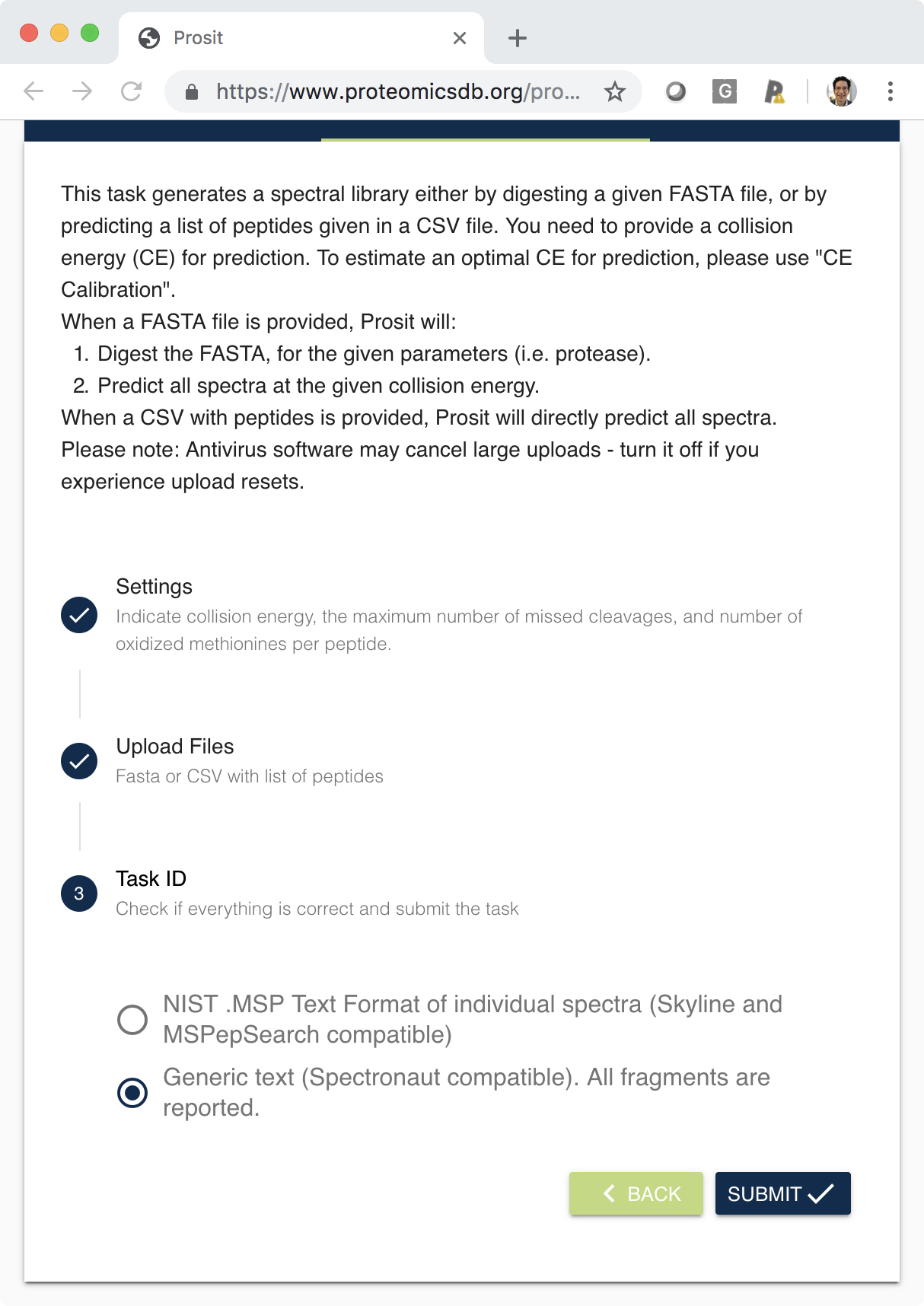

After the job has been submitted, record your Task ID. Depending on the size of the FASTA, your task may take hours to days to process. You can refresh the URL to check if your job has finished, or save the URL to check back at a later time. Once your task is complete, download the resulting files:

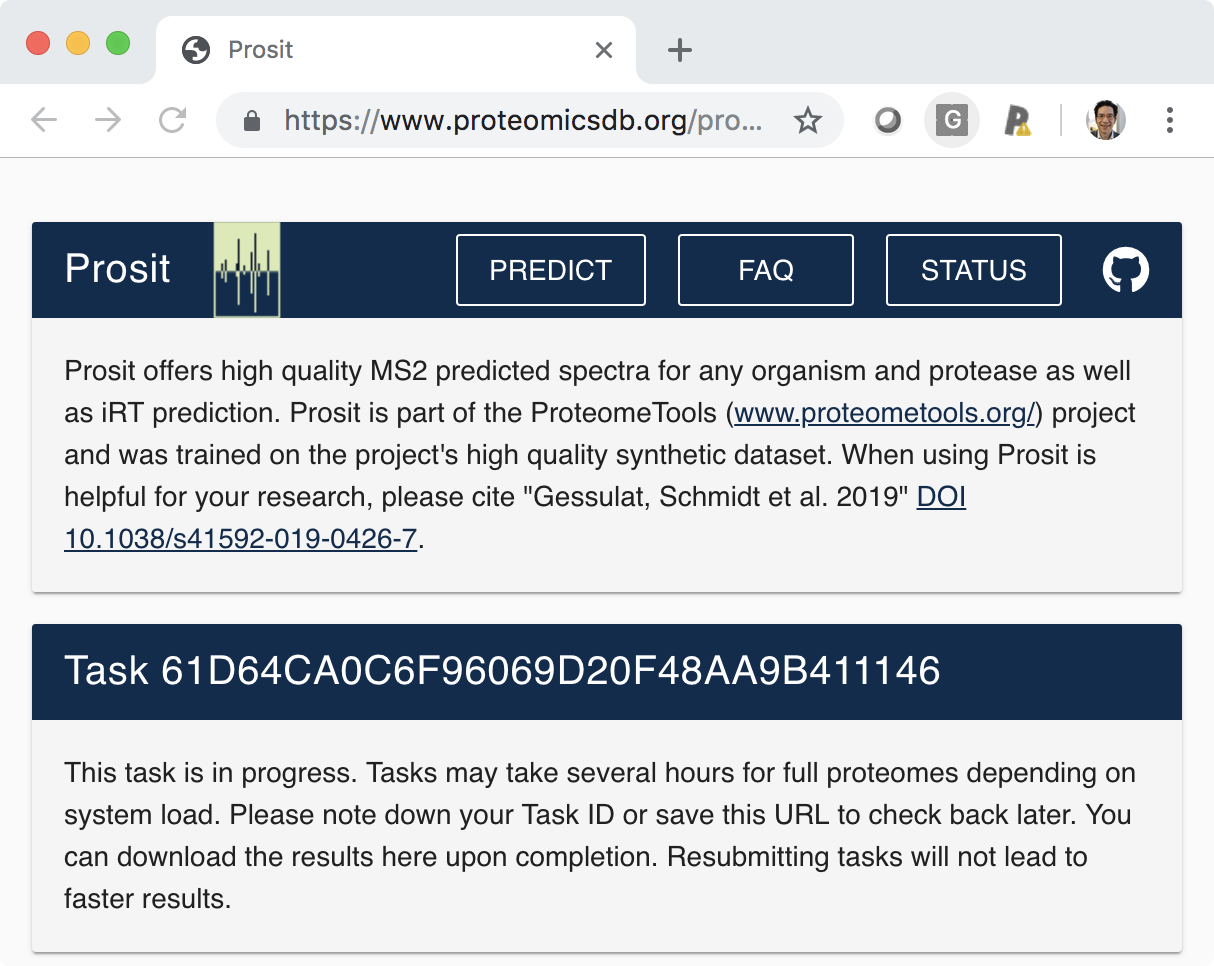

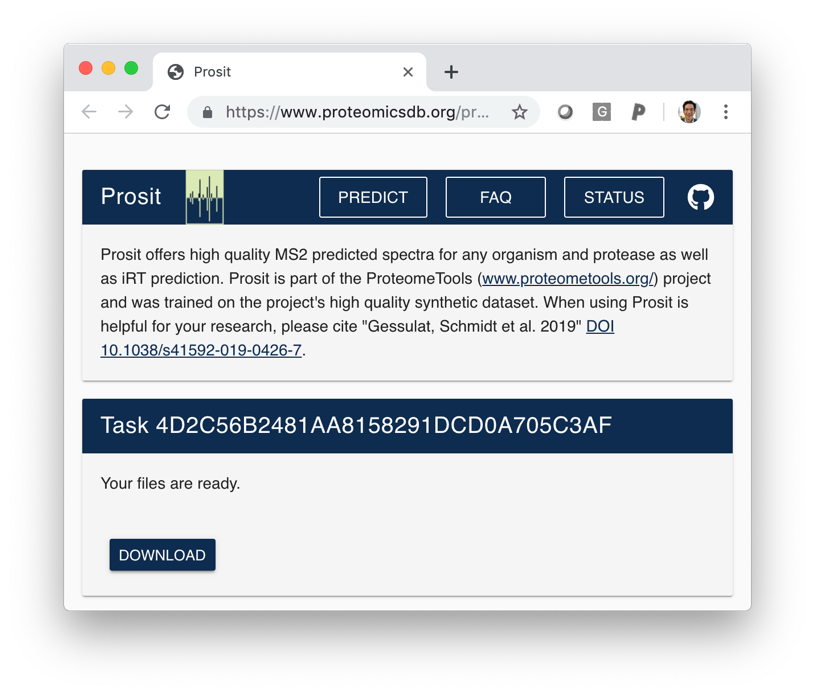

**Creating an EncyclopeDIA Library from Prosit CSV Output**

The “Convert/Create Prosit CSV from FASTA” menu option launches a dialog for building a predicted library. Upload the Prosit output CSV and your original FASTA database:

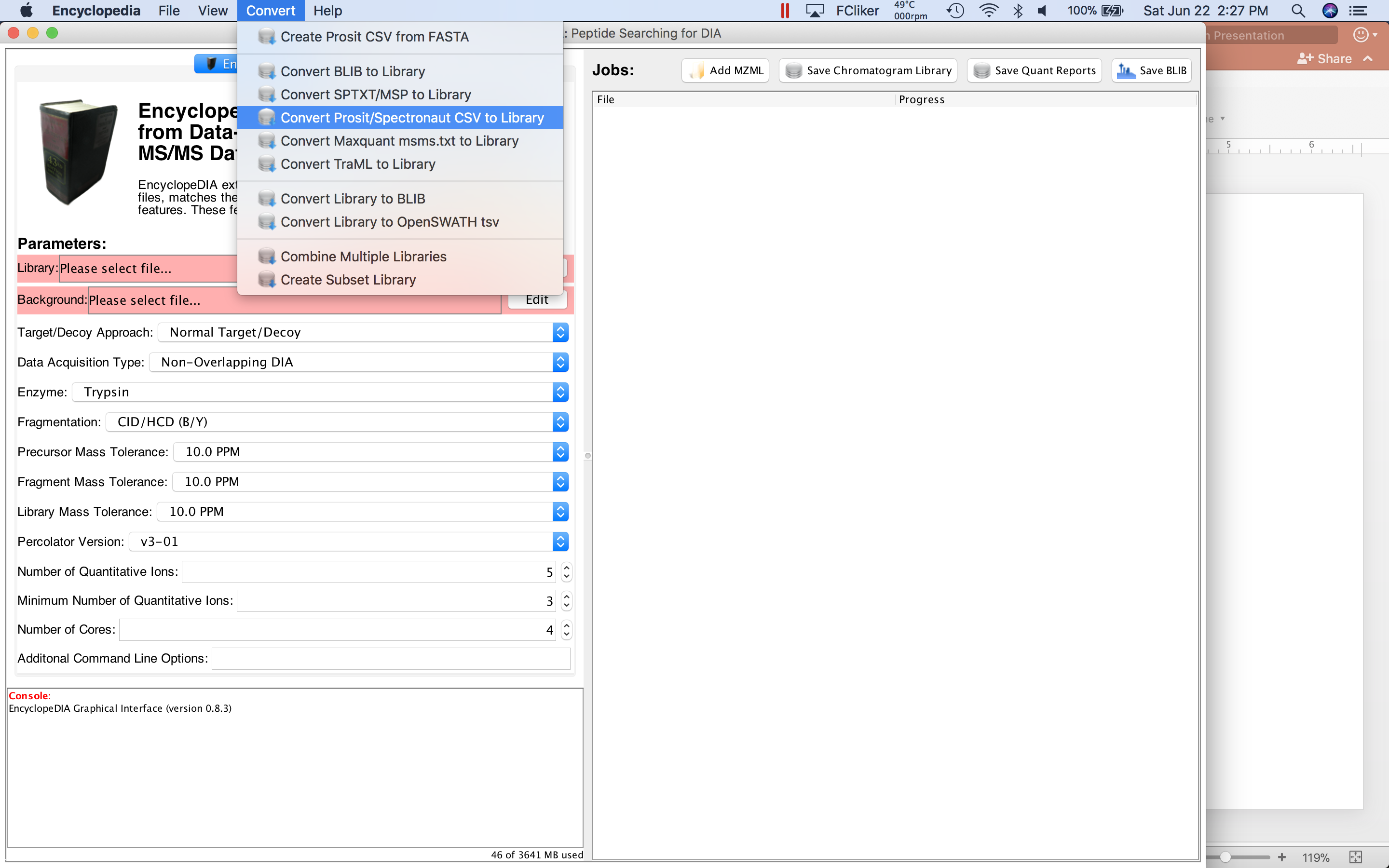

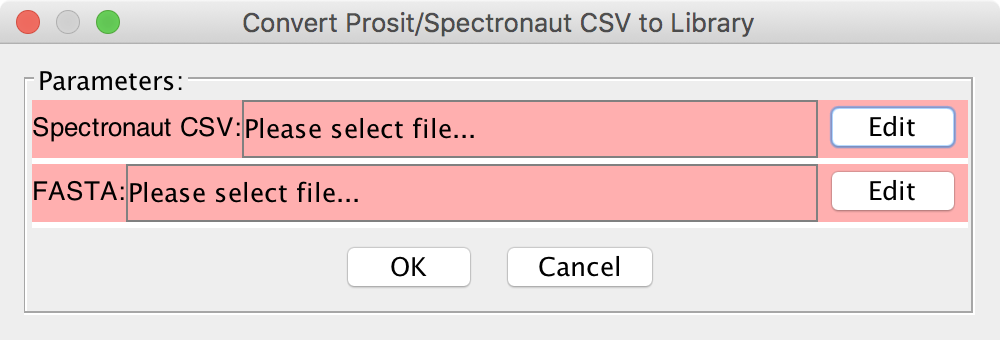

This will create a DLIB library in the same directory as your CSV.

**Making an Empirically-Corrected Library from Gas-Phase Fractionated Runs**

Using EncyclopeDIA parameter tab, specify the DLIB library and your FASTA database. Then set up your EncyclopeDIA search using settings appropriate for your experiment. For example, the following settings are appropriate for most orbitrap mass spectrometers:

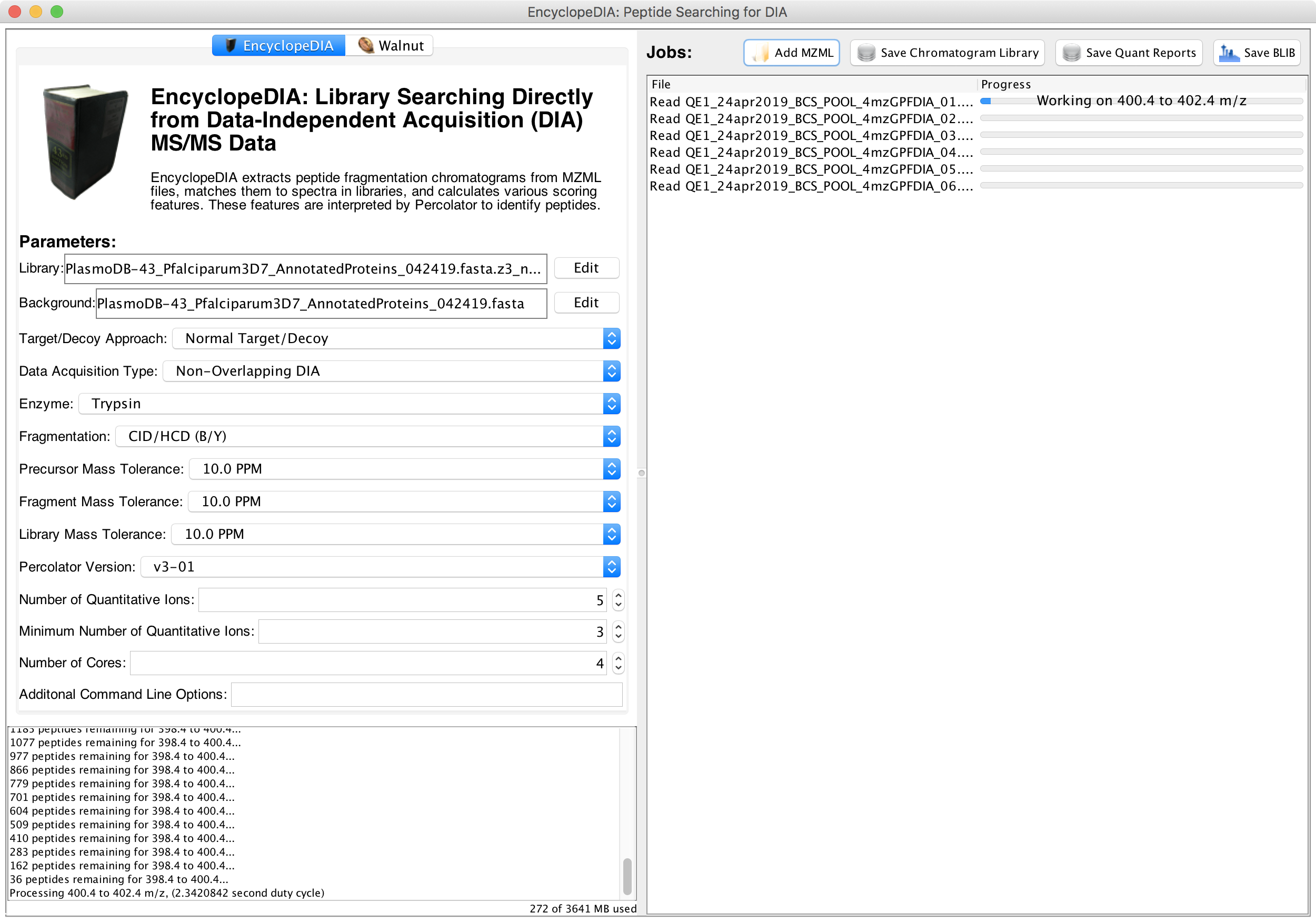

Then, queue up mzML files from your gas-phase fractionated runs. We recommend using MSConvert (Proteowizard) for building vendor-neutral mzML files from your vendor-specific raw files:

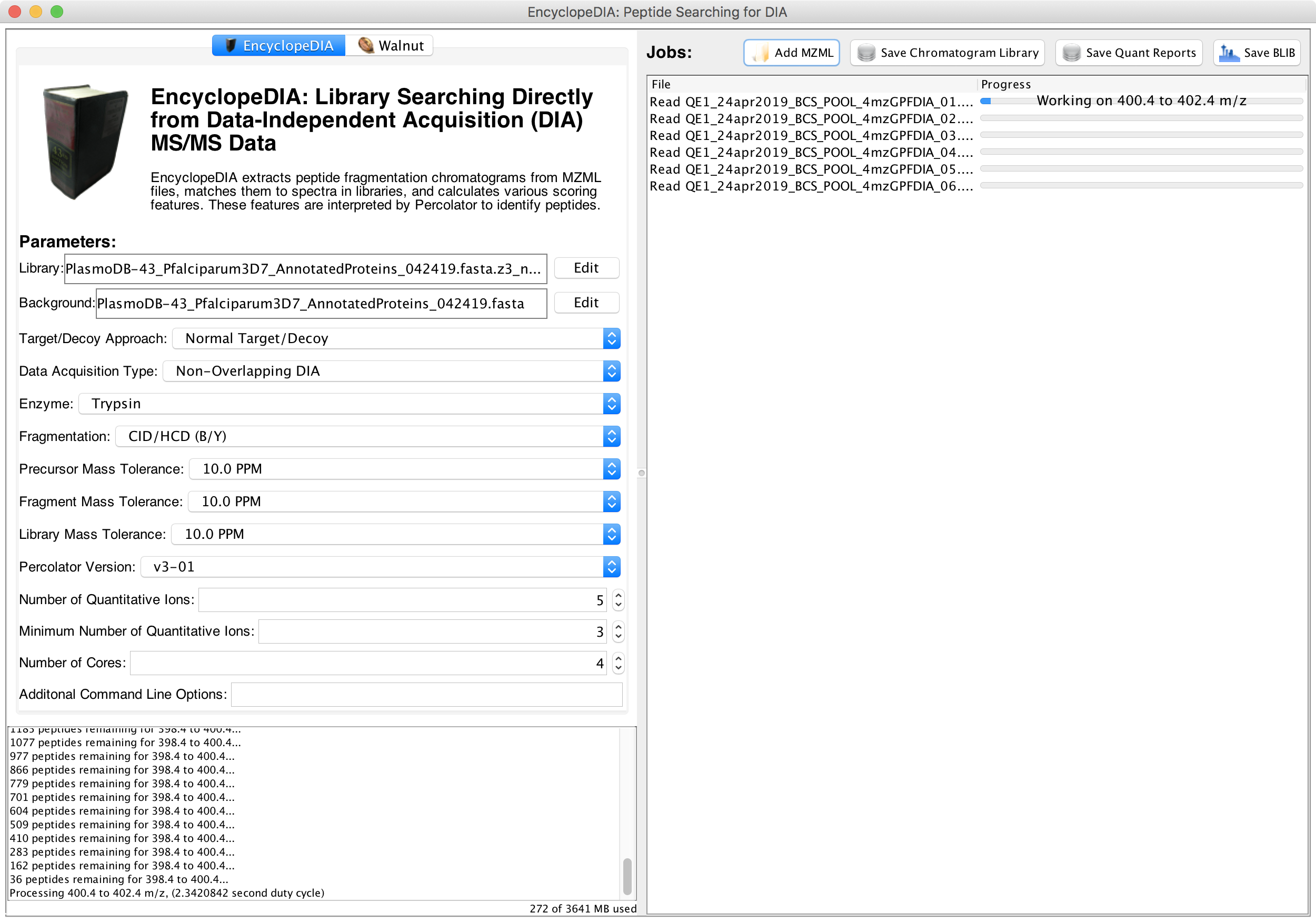

Once your files have finished processing, press the button “Save Chromatogram Library” to create your empirically-corrected library in an ELIB format. You can use this ELIB library in EncyclopeDIA, Skyline, or Scaffold DIA to analyze your single-injection DIA experiments.

**SUPPLEMENTARY FIGURES**

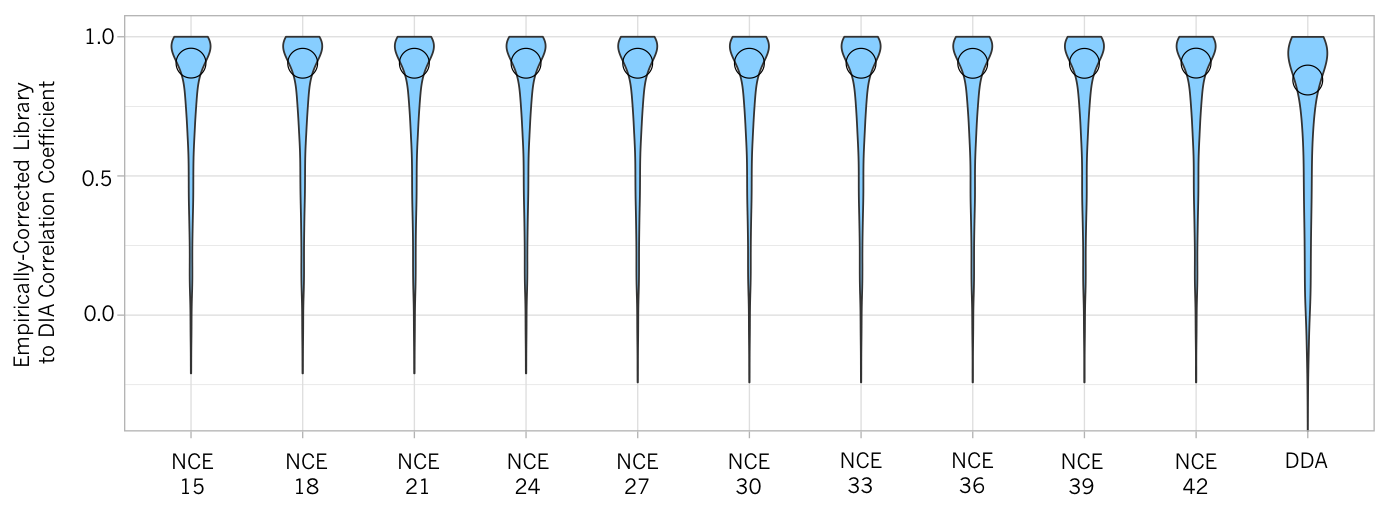

**Supplementary Figure 1**: **Comparison of empirically-corrected library and DDA library fragmentation.** Violin plots show that after empirical correction, the correlation between empirically-corrected library fragmentation and the integrated DIA chromatograms is consistently higher than the high-pH reverse-phase fractionated DDA spectrum library.

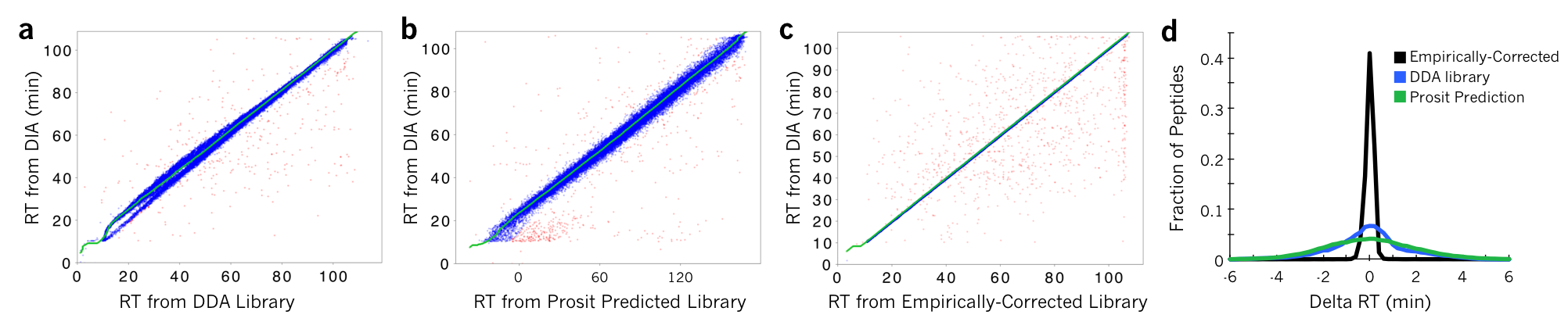

**Supplementary Figure 2**: **Comparison of empirically-corrected library and DDA library retention times.** Retention time (RT) alignment between one single-injection yeast replicate and (**a**) the DDA library (**b**) the Prosit predicted library, and (**c**) the resulting empirically-corrected library. Even though the DDA library was acquired on the same instrument with the same chromatography setup, there is error (including some minor streaking) between the fractionated and the single-injection retention times due to matrix effects. Green lines indicate the calculated non-linear retention time warping function. (**d**) Density curves showing the fraction of library peptides as a function of delta retention time after retention time warping. The empirically-corrected libraries have improved fragmentation and retention time accuracy over the predicted library and the DDA library, as GPF does not affect peptide interactions with matrix.

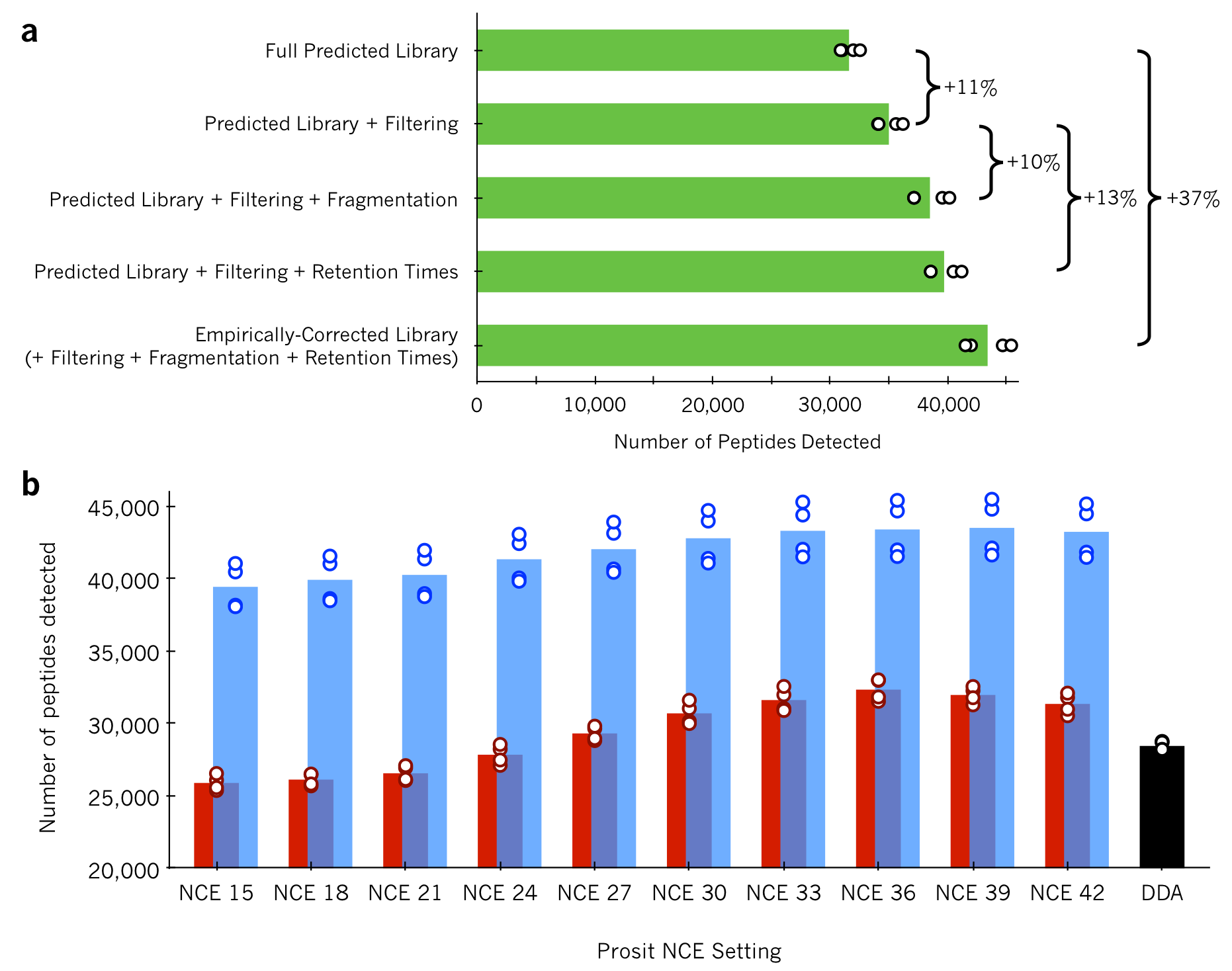

**Supplementary Figure 3**: **Additive effect of library size filtering, retention time, and fragmentation patterns.** (**a**) The number of yeast peptides detected at 1% peptide FDR in single-injection DIA runs (N=4) using the NCE=33 chromatogram library, where either the retention times or fragmentation patterns have been switched with the predicted Prosit values. Compared to the predicted spectrum library search, an 11% increase comes from simply using a narrowed peptide selection. DIA-based retention times and fragmentation patterns provide a 13% and 10% increase over this, respectively. Comparing the chromatogram library and the predicted library detections (37% increase), these percentage gains appear to be nearly multiplicative (i.e., 111%*110%*113%=138%), indicating that all 3 factors are independent and of roughly equal importance. (**b**) The same single-injection DIA runs (N=4) searched against either the Prosit predicted library (red bars) at various NCE settings, the resulting chromatogram libraries (blue bars), or a high-pH reverse-phase fractionated DDA spectrum library (black bar).

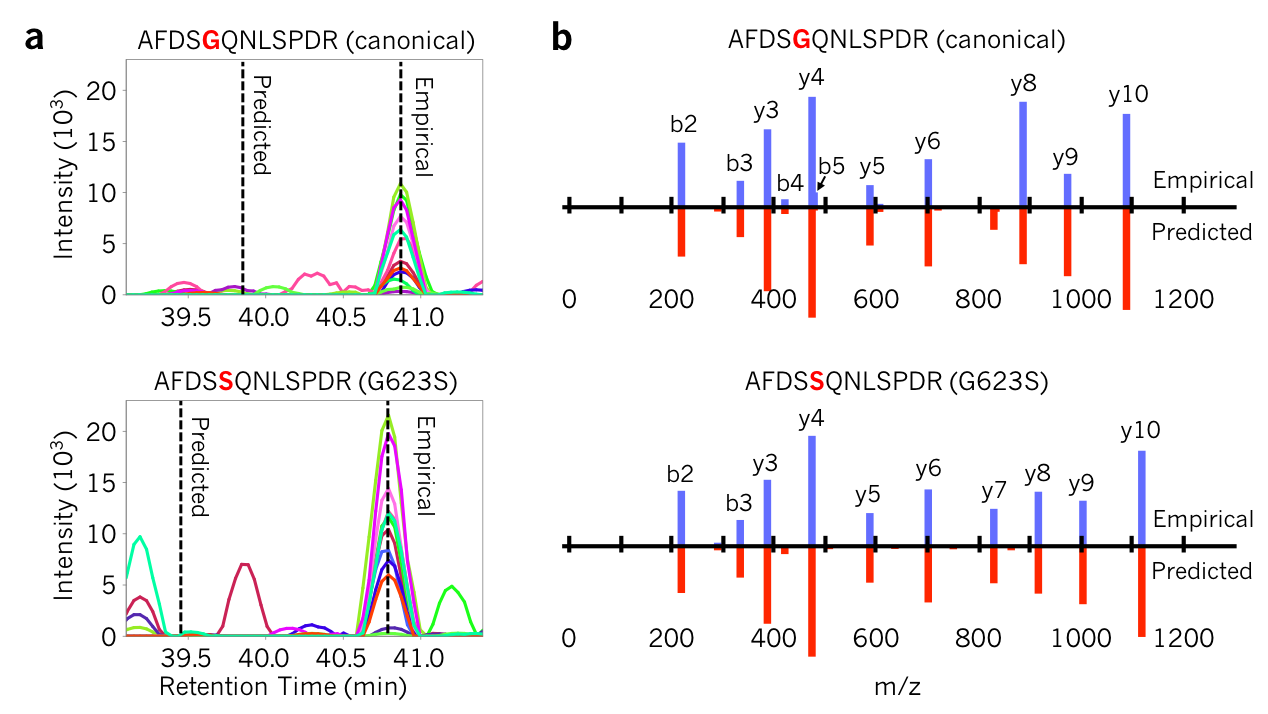

**Supplementary Figure 4: Heterozygous missense variants can produce difficult to separate peptides with DIA.** (**a**) Predicted and empirical retention times for the AFDSGQNLSPDR peptide and the G623S variant peptide from the kinase EEF2K show that these peptides elute at essentially the same time. (**b**) Fragmentation patterns for the AFDSGQNLSPDR peptide and the G623S variant peptide from the kinase EEF2K show that more than half of the detected fragment ions have the same m/z (y1-y7 and b1-b4). Relative fragmentation patterns are shown as butterfly plots with empirical intensities (blue) above predicted intensities (red).

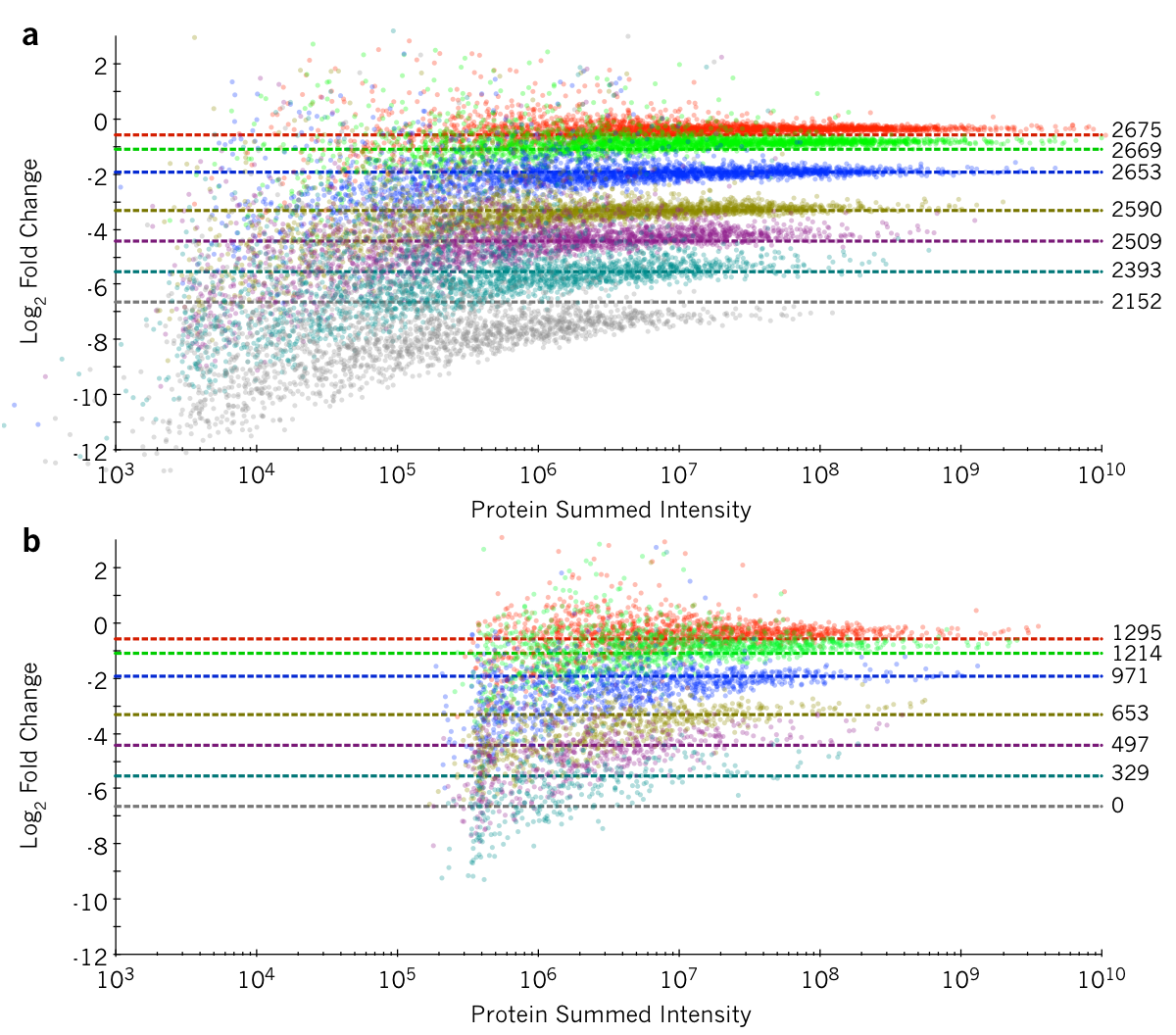

**Supplementary Figure 5**: **Quantification of *P. falciparum* proteins.** Quantitative ratios using either (**a**) DIA or (**b**) DDA for *P. falciparum* proteins at 7 different dilution ratios with red blood cell lysates (red=2:1, green=7:8, blue=4:15, gold=1:9, purple=2:41, cyan=2:91, and gray=1:99) relative to the protein intensity (summed from peptide intensities). Dashed lines indicate the expected ratio, where the number of proteins in each measurement batch are indicated at the right. No *P. falciparum* proteins were detected or quantified in the 1:99 dilution sample using DDA.

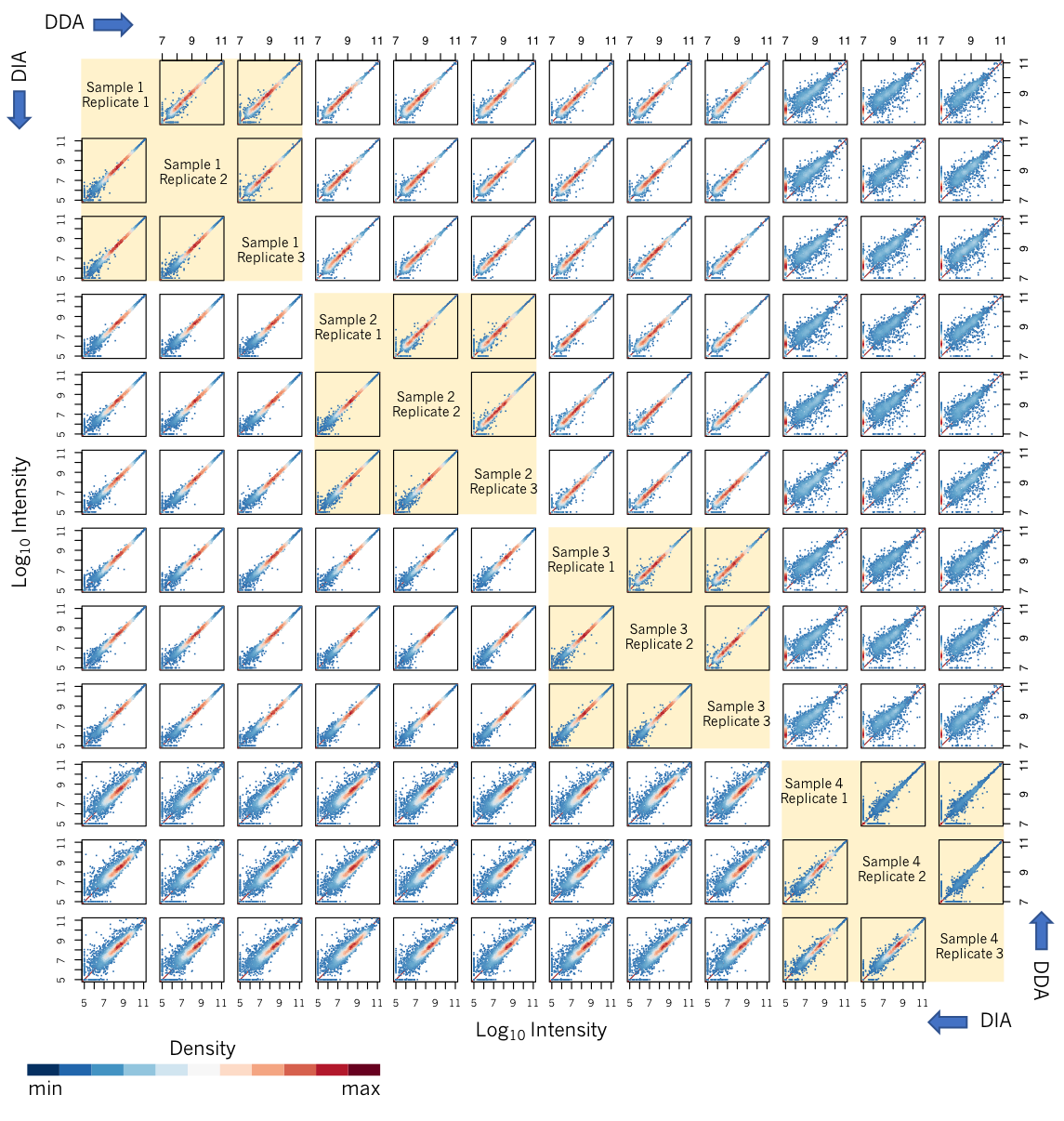

**Supplementary Figure 6**: **Quantitative robustness of single injection DIA and DDA.** Quantile-Quantile (Q-Q) plots for 4 culture replicates (each with 3 single-injection technical replicates) measured with DDA (upper right plots) and DIA (lower left plots). DDA quantitation was performed using MaxQuant (with “match between runs” enabled) and DIA quantitation was performed using EncyclopeDIA. Samples 1-3 were highly enriched for *P. falciparum* via magnetic-activated cell sorting, and quantitative consistency between both culture and technical replicate DDA runs and replicate DIA runs is very high. Based on increased human protein detection, Sample 4 appeared to contain some concentration of red blood cells. In DIA this resulted in some additional measurement scatter in the Q-Q plots; however, the majority of proteins can still be quantified accurately. On the other hand, in DDA the majority of quantitative measurements are missing in Sample 4, as indicated by the red densities against the y-axis of the Q-Q plots.
